# Source-Sink Modifications Affect Leaf Senescence and Grain Mass in Wheat

**DOI:** 10.1101/647743

**Authors:** Xuemei Lv, Yan Zhang, Yunxiu Zhang, Shoujin Fan, Lingan Kong

**Author notes:** **Corresponding author:** Lingan Kong, Ph.D., College of Life Science, Shandong Normal University, 88 Wenhuadong Road, Jinan City 250014, China.

## Abstract

A field experiment was performed in wheat to investigate the responses of flag leaf and grain to sink/source manipulations. The results showed that half-degraining delayed but defoliation (only flag leaf left) enhanced the leaf senescence. Sink/source manipulations influenced the content of reactive oxygen species of flag leaf and the content of phytohormones including cytokinins, indoleacetic 3-acid, gibberellin 3, salicylic acid and jasmonic acid in the defoliated flag leaf (DL) and grain (DG), half-degrained flag leaf (HL) and grain (HG). An iTRAQ based quantitative proteomic analysis indicated that at 16 days after manipulation a total of 97 and 59 differentially expressed proteins (DEPs) from various functional categories were observed in HL and DL groups, respectively, compared with control and 115 and 121 DEPs were observed in HG and DG groups, respectively. GO annotation terms of DEPs mainly included carbon fixation, hydrogen peroxide catabolic process, chloroplast and cytoplasm, oxidoreductase activity and glutamate synthase activity in flag leaf of manipulated plants; organonitrogen compound metabolic process, cytoplasm, vacuolar membrane, CoA carboxylase activity, starch synthase activity and nutrient reservoir activity in grain of manipulated plants. KEGG pathway enrichment analysis revealed that photosynthesis, carbon, nitrogen and pyruvate metabolisms and glycolysis/gluconeogenesis were the most remarkable processes for sink/source manipulations. Sink/source manipulations affected the activities of α- and β- amylases and proteinases. Ultimately, manipulations changed the mass per grain. In conclusion, manipulations to change the sink/source ratio affect the levels of hormones, activities of hydrolytic enzymes, metabolisms of carbon, nitrogen and other main compounds, stress resistance, the leaf senescence, and ultimately influence the grain mass.

## Introduction

The grain yield of cereals is determined by the synergistic reaction between source activity and sink capacity. Source supply reserves accumulated there to sink (the developing grains). Sink strength is determined by the number and potential size of grains per stem, depending on the capacity to actively obtain from photosynthetic assimilates and reserves in the vegetative organs and accumulate these compounds.

Wheat (*Triticum aestivum* L.) productivity is generally considered to be a sink-limited under favorable conditions with grain development regulated by the assimilating capacity during grain filling, is hardly limited by the source (Foulkes *et al*., 2011), because that the wheat source generally has the capacity to provide adequate assimilates to the developing grains. However, the inconsistent conclusions have also observed. An increase in the source/sink ratio do not affect the grain mass (Calderini and Reynolds 2000; Calderini *et al*., 2006). Defoliation in old germplasm of bread wheat does not cause source limitation to grain filling (Kruk *et al*., 1997). The grain yields are co-limited both by the source and the sink in wheat (Kruk *et al*., 1997; Beed *et al*., 2007). Experiments with manipulation of assimilate availability during grain filling found that the wheat yields are mainly limited by the sink size (Borŕas *et al*., 2004). Slafer and Savin (1994) reported that the grain yield of wheat is either sink-limited or co-limited by both source and sink but never source-limited. Therefore, better understanding of source-sink interaction would help propose strategies to improve yield potential using genetic avenues.

A number of phytohormones play vital roles in regulating leaf senescence and grain filling process in crops as signaling molecules. phytohormones play defernet roles in plant senescence, such as ABA and jasmonic acid (JA) inducing senescence, while cytokinins (CKs), auxin, and gibberellin (GA) show a contrasting response (Yang *et al*., 2000; Chang and Zhu, 2017; Jan *et al*., 2019). In addition, various studies have outlined the cross-talk among various hormones. They can be directly involved in the regulation of senescence or function antagonistically. In addition, interactions of phytohormones with other factors, such as nitrogen status and sugar signaling, play a critical role in regulating source and sink communication between (Paul and Foyer, 2001; Thomas and Ougham, 2014). Therefore, the regulation of senescence by hormones is a complicated process that needs deeper investigation.

As a powerful technique to perform quantitative proteome analysis, isobaric tag for relative and absolute quantitation (iTRAQ) allows identification of more numerous proteins and can provide more reliable quantitative information than traditional 2-DE analysis (Karp *et al*., 2010). With a sufficient number of proteins it is possible to construct pathways and perform protein-protein interaction (PPI) analyses (Wang and You, 2012). The proteomics studies in wheat using iTRAQ were primarily on performed for investigating the protein responses to stress such as drought, reactive oxygen species (ROS) stress and nutrient deficiency to assess the effects of environmental factors on profile characteristics and metabolic pathway (Ford *et al*., 2011; Wang and You, 2012). To date, however, quantitative proteomics studies based on iTRAQ analysis on wheat senescence and grain development when subjected to sink-source manipulations have not been reported.

Modification of the source-sink relationship by changing the levels of competition among developing grains and/or the the assimilate availability has been previously used as manipulations after grain setting to understand the effects of source/sink balance on grain development, aiming at suggesting strategies to increase the grain yield of wheat (Serrago *et al*., 2013; Slafer and Savin, 1994; Ma *et al*., 1996) and barley (Cartelle *et al*., 2006). Unfortunately, although much effort has been made, we are still far from fully understanding source-sink interaction and even further from rational manipulation of the source-sink relationship (Chang and Zhu, 2017). How it can be regulated by cultivation means and genetic manipulations remained unclear.

The study reported here was designed to examine the impact of manipulating the source/sink ratio on the physiological modifications and protein expression in flag leaf and grain, aiming to determine how the leaf senescence and grain yield is regulated by changing the availability of potential assimilates for per grain. These results might promote better understanding of the roles of source-sink relationship in grain development and finding effective avenues, such as genetic modifications in breeding, to further increase cereal grain yield.

## Results

### NDVI, PRI and SPAD values

The changes in NDVI, PRI and SPAD values of the flag leaves exhibited similar trends (Table 1). Three parameters were gradually decreased with the progress of grain filling. The PRI values showed the greater sensitivity to sink-source modifications than SPAD and NDVI. Significant differences in PRI values were observed between three treatments and between growth stages. However, the differences in NDVI values were only observed between half-degraining and defoliation at 16 and 24 Days after manipulation (DAM). At 16 DAM, significant difference in SPAD values was only observed between half-degraining and defoliation. At 24 DAM, the highest level was observed in half-degrained plants, followed by significant decrease in control and further in defoliated plants (Table 1).

**Table 1.** Effects of sink-source manipulations on the PRI, NDVI and SPAD values of flag leaf.

| DAT |  | PRI | NDVI | SPAD |
| --- | --- | --- | --- | --- |
| 8 | Control | 0.030b | 7.98ab | 58.77ab |
|  | Half-degraining | 0.032a | 8.14a | 59.83a |
|  | Defoliation | 0.028b | 7.89ab | 58.07ab |
| 16 | Control | 0.026cd | 7.47cd | 57.27bc |
|  | Half-degraining | 0.028bc | 7.74bc | 58.03ab |
|  | Defoliation | 0.024e | 7.26d | 56.03c |
| 24 | Control | 0.021f | 6.21f | 38.43e |
|  | Half-degraining | 0.025de | 6.57e | 40.73d |
|  | Defoliation | 0.018g | 6.14f | 33.17f |

### Chlorophyll fluorescence

The value of flag leaf *Fv/Fm* remained unaffected during experiment period for half-degraining treatment compared with the intact control, while the value was significantly lower for defoliation than control (Fig. 1). The difference in Φ_PSII_ value between half-degraining and control was only observed at 16 DAM, while the Φ_PSII_ value were significantly lower for defoliation than degrained and control plants (Fig. 1).

**Fig. 1.**
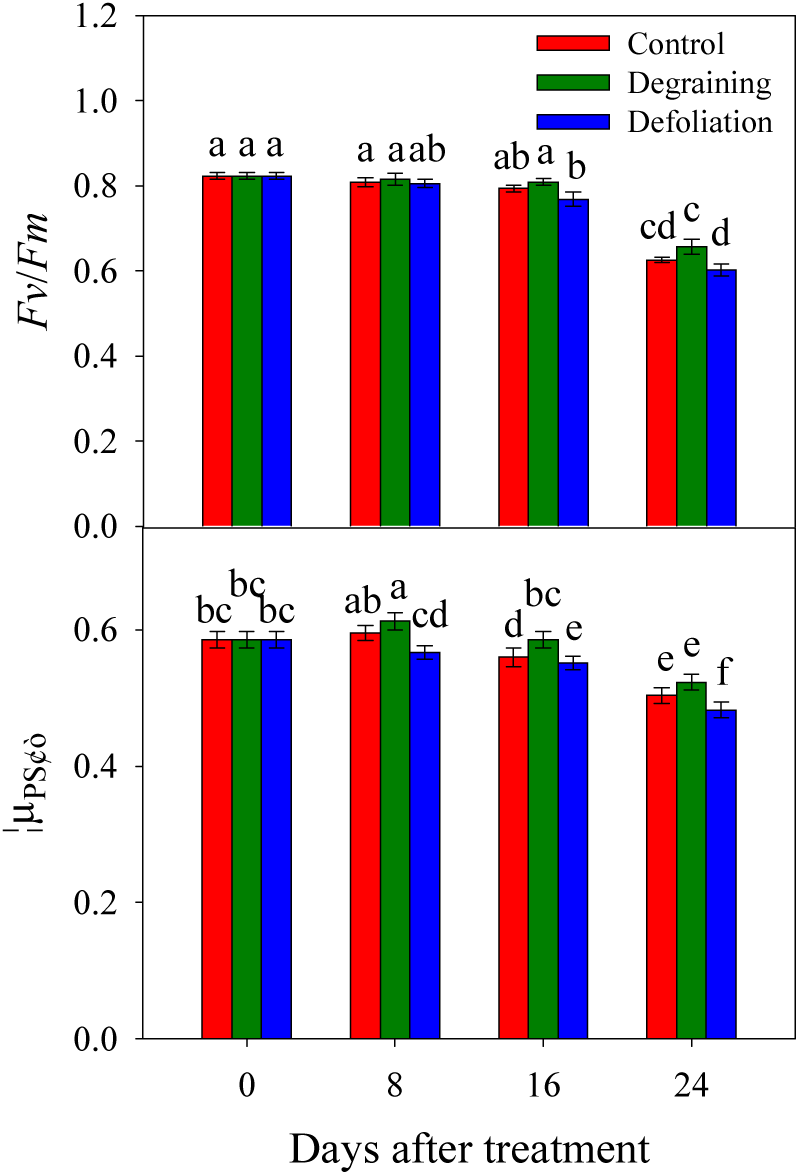
Effects of sink/source manipulation on fluorescence parameters of flag leaves in wheat. (A) *Fv/Fm*: maximal efficiency of PSII photochemistry; Φ_PSII_: actual PSII efficiency. Each value is the mean ± standard deviation (SD) from at least six leaves. The columns labeled with different letters differed significantly at *p<*0.05 according to Duncan’s multiple range test.

### Chloroplast Structure

At 16 DAM, visual observation did not find difference in senescing flag leaf between three treatments, however, the obvious difference in cell ultrastructure of leaf tissues was observed under transmission electron microscopy (Fig. 2). The chloroplasts were spherical in all three treatments. In cells of degrained flag leaf (Fig. 2A), the chloroplasts contained a larger number of thylakoids and the matrix of the chloroplasts was still dense in comparison to the control (Fig. 2B) and defoliation treatment (Fig. 2C). In cells of defoliated flag leaf, numerous small vesicles or membrane-like fragments were observed in the cytoplasm (Fig. 2C). The membranes constituting the thylakoids were more distinct and abundant in cells of degrained flag leaf. But in control and defoliation treatments, the structure of the thylakoids was characterized by loss of the parallel arrangement of the grana lamellae in some chloroplasts, and some of the thylakoids became swollen. In cells of flag leaf with defoliation treatment, the number of chloroplasts decreased noticeably.

**Fig. 2.**
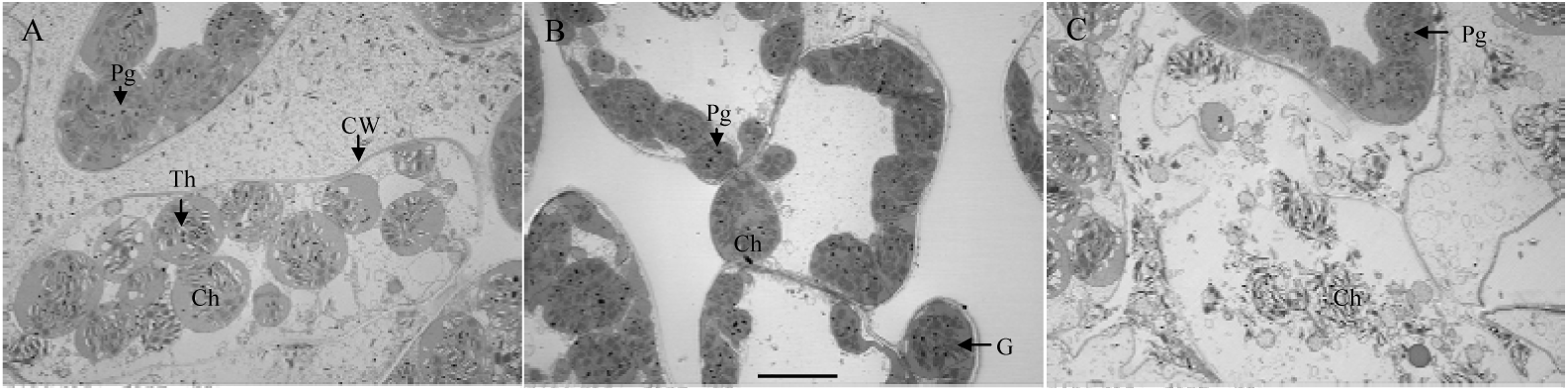
Transmission electron micrographs showing the ultrastructure of the chloroplast at 16 DAM in flag leaves of control (A), half-degraining (B) and defoliation (C) treatments. Bars: 2 μm; Ch, chloroplast; CW, cell wall; G, granum; Mt, mitochondrion; Pg, plastoglobuli; Th, thylakoid.

### Antioxidant Enzyme Activity of Flag Leaf

At 8 DAM, no significant difference in superoxide dismutase (SOD) activity of flag leaf was observed between half-degraining, defoliation and control. At 16 and 24 DAM, half-degraining significantly decreased the SOD activity compared with the control (*p*<0.05). However, defoliation increased the SOD activity at 16 (*p* > 0.05) and 24 (*p*<0.05) DAM (Fig. 3a). The peroxidase (POD) activity exhibited the almost identical changing trend as SOD activity during the period of 8 to 24 DAM (Fig. 3b). At 8 and 24 DAM, half-degraining significantly decreased the catalase (CAT) activity with respect to the control (*p*<0.05). The defoliation treatment decreased the CAT activity at 8 DAM (*p*<0.05), but caused sharp increases in the activity thereafter and thus induced significantly enhancement of CAT activity 24 DAM (*p*<0.05) (Fig. 3c). Even though the higher activities of these enzymes, especially at 24 DAM, the content of ROS was still higher in flag leaf of defoliated plants. However, no difference in ROS content was observed in flag leaf between half-degraining and intact plants (Fig. 3d).

**Fig. 3.**
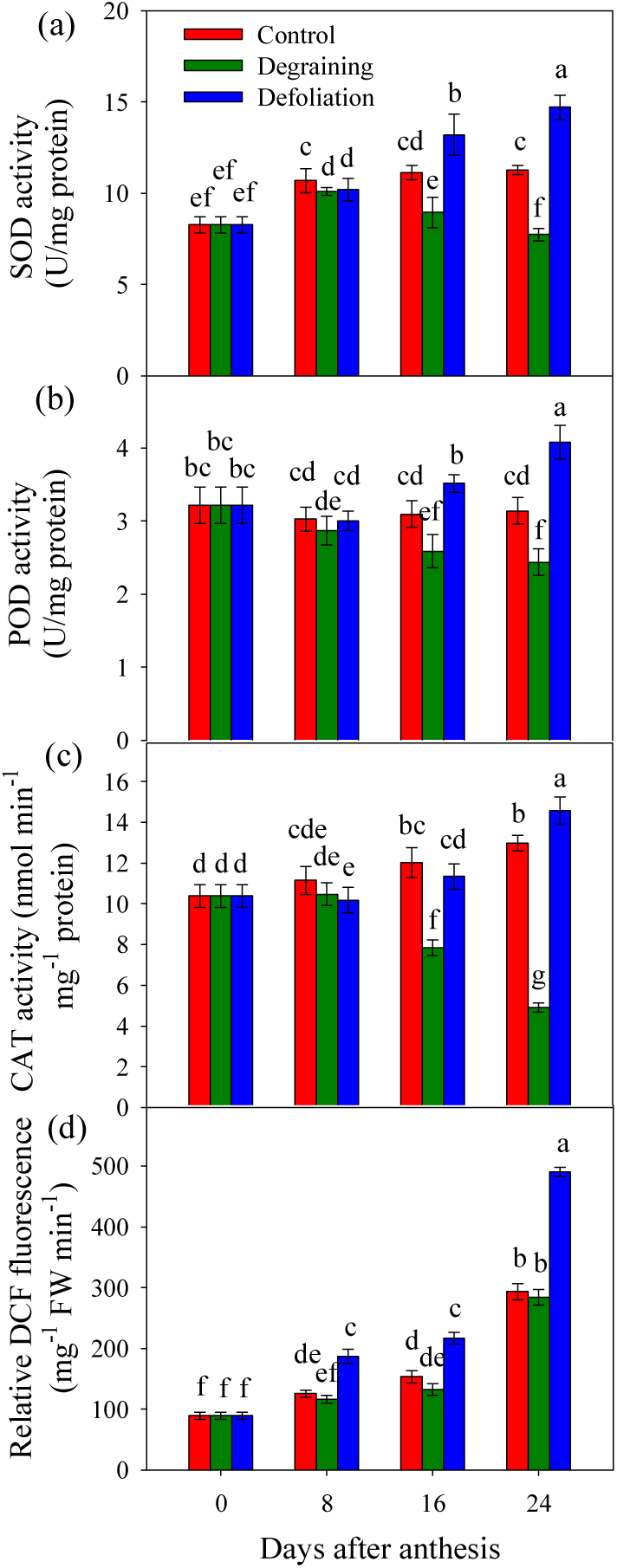
Effects of sink-source manipulations on antioxidant enzyme activities and the relative DCFH-DA fluorescence intensity of wheat flag leaves. Results are mean ± SD.

### Effect of Sink-Source Manipulations on Endogenous Hormones in Flag Leaves and Grains

The total content of zeatin + zeatin ribuside + kinetin was used to evaluate the changes of CTKs after manipulation. Both half-degraining and defoliation treatments decreased the CTKs content in flag leaf at 8 (*p*<0.05), 16 (*p*<0.05) and 24 (*p*>0.05) DAM. However, the CTKs content decreased in the remained grain of half-degrained plants at 8, 16 and 24 (*p*<0.05); whereas it significantly increased for defoliation at three time points. Interestingly, the ratio of grain to leaf CTK contents was significantly decreased for half-degraining while significantly increased for defoliation (Table 2).

**Table 2.**
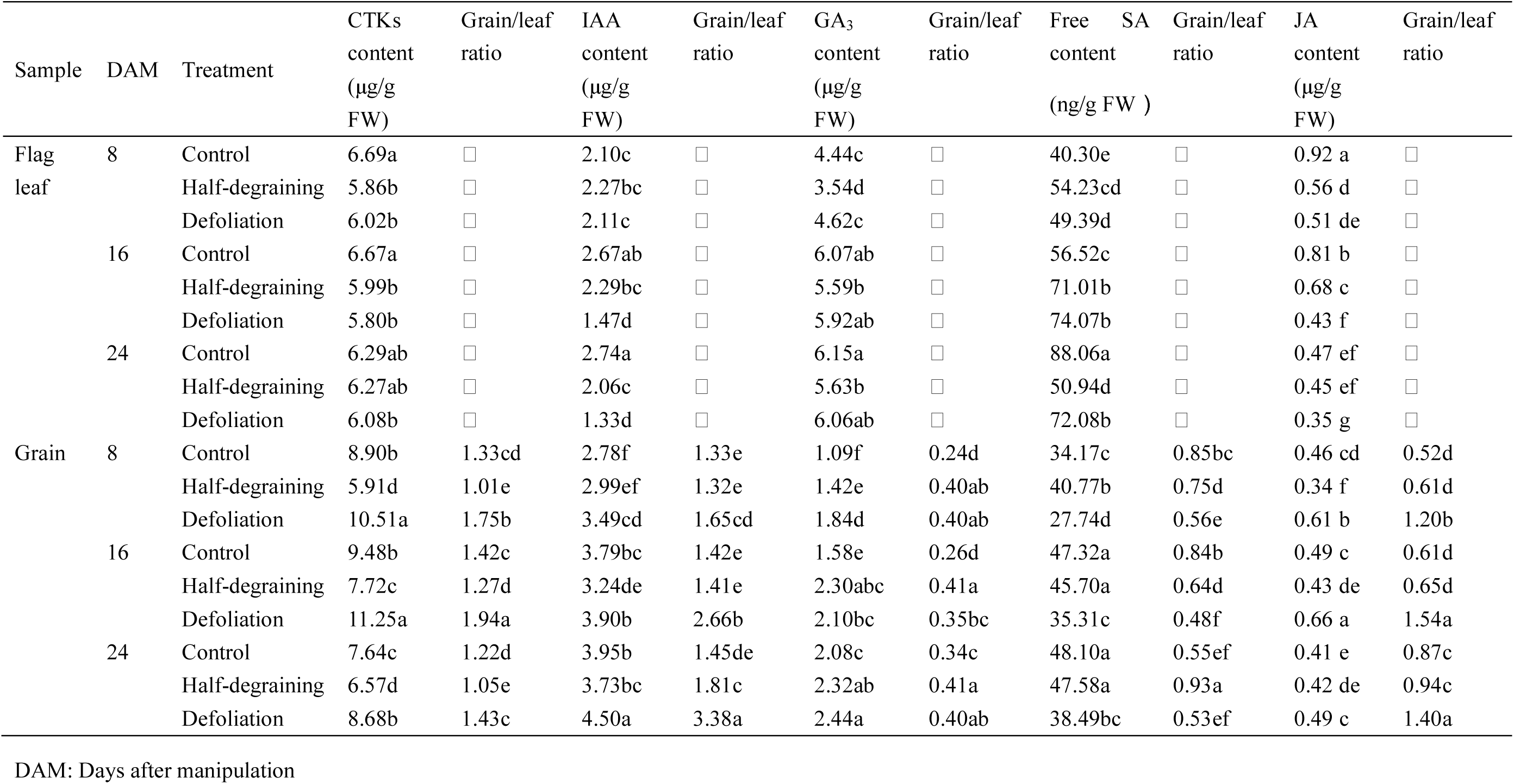
Effects of sink and source manupulations on the phytohormone contents in wheat flag leaves and grains.

No significant difference in flag leaf indoleacetic 3-acid (IAA) content was observed between three treatments at 8 DAM. At 16 and 24 DAM, the highest level of the flag leaf IAA content was observed in control, followed by significant decreases in half-degraining (*p*<0.05) and further in defoliation (*p*<0.05) (Table 2). In grain, the IAA content increased for defoliation treatment at 8, 16 and 24 DAM (*p*<0.05), while no difference was observed between control and half-degraining treatment. As a result, the ratio of grain to leaf IAA contents was significantly increased for half-degraining at 24 DAM and for defoliation at three time points (Table 2). In flag leaf, half-degraining significantly decreased the GA_3_ content at 8, 16 and 24 DAM, but no difference was observed between defoliation and control. In grain, both manipulations increased the GA_3_ content at 8, 16 and 24 DAM. Thus, both manipulations increased the ratio of grain to leaf IAA content (Table 2).

Half-degraining significantly increased the free salicylic acid (SA) content in flag leaf at 8 and 16 DAM and in grain at 8 DAM. Defoliation increased the free SA content in flag leaf during the experimental period but had the reverse effects on grain. As a consequence, the ratio of grain to leaf SA content was significantly decreased for both manipulations decreased at 8 and 16 DAM, while the ratio was significantly increased for half-degraining at 24 DAM (Table 2). In flag leaf, both manipulations decreased the JA content throughout the experimental time; in grain, half-degraining reduced the JA content at 8 and 16 DAM, while defoliation significantly enhanced the JA content throughout the experimental time, resulting significantly increases in the ratio of grain to leaf JA content for defoliation at three time points but similar between control and half-degraining manipulation (Table 2).

### Proteome Profiles of Wheat Flag Leaves and Grains under Different Sink-Source Modifications

To investigate the proteome alterations of wheat flag leaves and grains due to different sink-source modifications, iTRAQ-based quantitative proteomics analysis was conducted. Totals of the 2821 proteins in flag leaves and 2467 proteins in grains were identified reliably at a global FDR of 1%. In flag leaves of half-degrained plants, 97 proteins were assigned as differentially expressed compared with intact plants, including 80 increased proteins and 17 decreased proteins. In flag leaves of defoliated plants, 59 proteins were assigned as differentially expressed compared with intact plants, including 54 increased proteins and 5 decreased proteins. In grains of half-degrained plants, 115 proteins were assigned as differentially expressed compared with intact plants, including 43 increased proteins and 72 decreased proteins. In defoliated plants, 121 proteins were assigned as differentially expre**s**sed compared with intact plants, including 47 increased proteins and 74 decreased proteins (Fig. 4; Table S1).

**Fig. 4.**
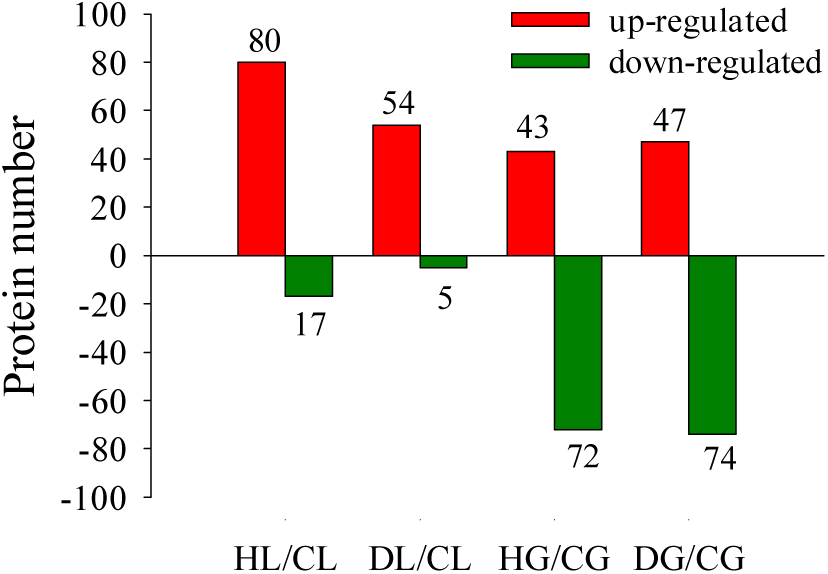
Number of differential expressed proteins (DEPs) of different groups. HL, half-degrained leaf; DL, defoliation leaf; CL, control leaf; HG, half-degrained grain; DG, defoliation grain; CG, control grain.

### Gene Ontology (GO) Analysis

To illustrate the major responsive processes in wheat flag leaves and grains, enrichment analysis was performed on the GO ontology. The top 20 of the GO terms to them the DEPs clustered in the biological process (BP), cell component (CC), and molecular function (MF) categories by GO analysis, respectively, is shown in Fig. 5.

**Fig. 5.**
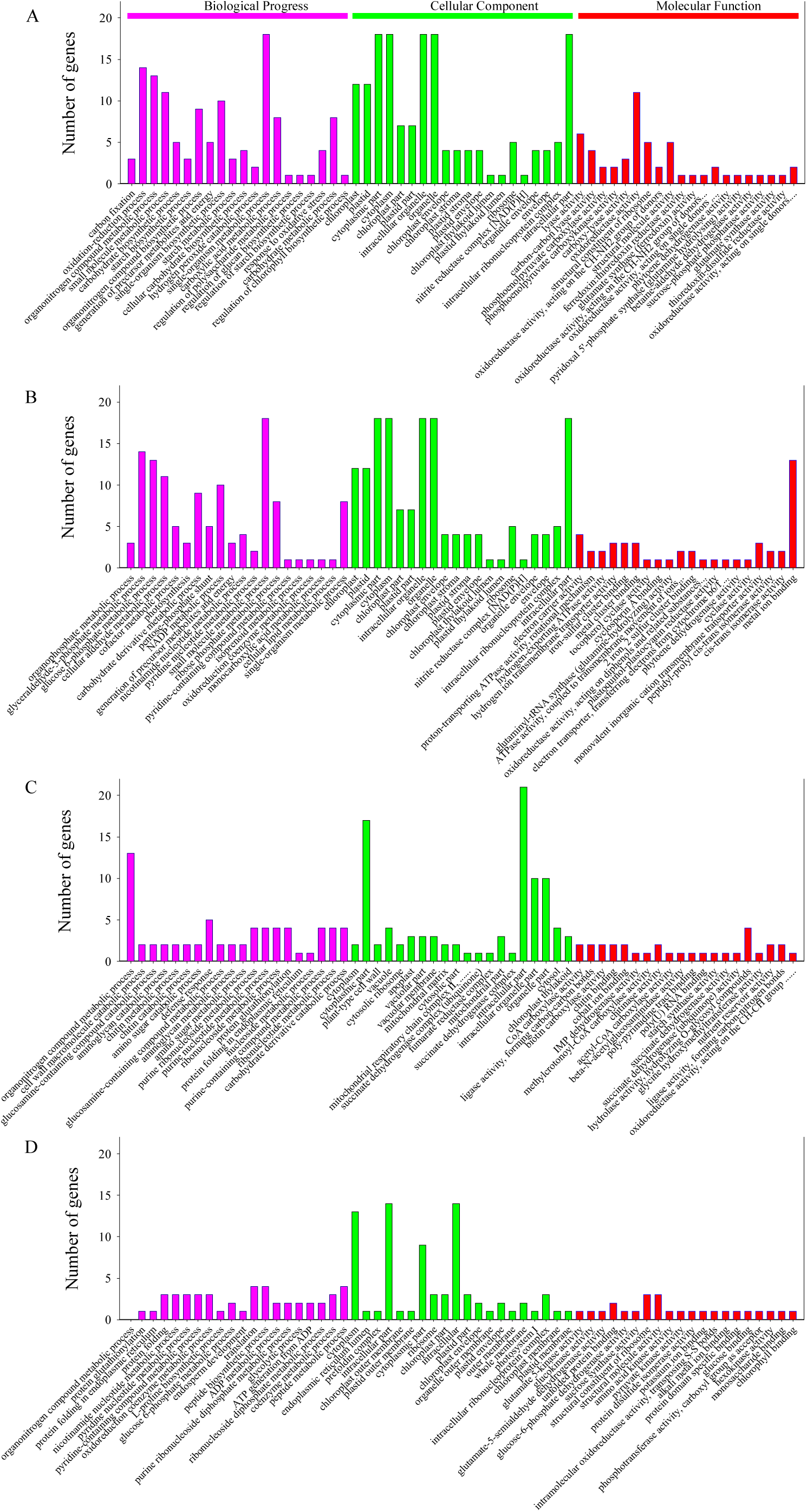
GO annotation of DEPs in manipulated flag leaf and grain compared to control in biological process, cellular component and molecular function. (A), Flag leaf: half-degraining vs control; (B), flag leaf: defoliation vs control; (C), grain: half-degraining vs control; (D), grain: defoliation vs control.

In flag leaf, the BP analysis revealed that the DEPs identified in comparison of HL with CL were mainly classified into carbon fixation, oxidation-reduction process, organonitrogen compound metabolic process, carbohydrate biosynthetic process, starch biosynthetic process and etc. In MF analysis, the DEPs were classified into the organophosphate metabolic process, glyceraldehyde-3-phosphate metabolic process, glyceraldehyde-3-phosphate metabolic process, glucose 6-phosphate metabolic process, photosynthesis, carbohydrate derivative metabolic process, pentose-phosphate shunt, NADP metabolic process, cellular lipid metabolic process, single-organism metabolic process and etc. According to the CC analysis, half-degraining mainly affected the chloroplast, plastid, cytoplasmic part, cytoplasm, chloroplast part, plastid part, intracellular organelle, chloroplast envelope and chloroplast stroma (Fig. 5A; Table S2).

The BP analysis revealed that the DEPs identified in comparison of DL with CL were mainly classified into organophosphate metabolic process, glyceraldehyde-3-phosphate metabolic process, glucose 6-phosphate metabolic process, cellular aldehyde metabolic process, photosynthesis, carbohydrate derivative metabolic process and etc. In the MF analysis, responses to the half-degraining mainly included carbon-carbon lyase activity, phosphoenolpyruvate carboxylase activity, phosphoenolpyruvate carboxykinase activity, carboxy-lyase activity, oxidoreductase activity, ferredoxin:thioredoxin reductase activity and glutamate synthase (ferredoxin) activity. Responses to defoliation mainly included electron carrier activity, proton-transporting ATPase activity, hydrogen-exporting ATPase activity, hydrogen ion transmembrane transporter activity, iron-sulfur cluster binding, tocopherol cyclase activity and glutaminyl-tRNA synthase (glutamine-hydrolyzing) activity. Defoliation mainly affected chloroplast, plastid, chloroplast part, plastid part, thylakoid part, thylakoid membrane, chloroplast envelope, organelle subcompartment and photosystem II (Fig. 5B; Table S2).

In grain, the BP analysis revealed that the proteins identified in HG compared with CG were mainly classified into organonitrogen compound metabolic process, cell wall macromolecule catabolic process, amino sugar catabolic process, defense response, aminoglycan metabolic process, amino sugar metabolic process, protein folding in endoplasmic reticulum and nucleoside metabolic process. In comparison of DG/CG, the majority of identified proteins were classified into CoA carboxylase activity, ligase activity, forming carbon-carbon bonds, biotin carboxylase activity, methylcrotonoyl-CoA carboxylase activity, acetyl-CoA carboxylase activity, starch synthase activity, succinate dehydrogenase activity and etc. According to the CC analysis, half-degraining mainly affected cytoplasm, cytoplasmic part, vacuole, apoplast, vacuolar membrane, mitochondrial part, intracellular par and chloroplast thylakoid (Fig. 5C; Table S2).

For BP analysis, grain responses to defoliation mainly included organonitrogen compound metabolic process, protein glutathionylation, protein folding in endoplasmic reticulum, pyridine nucleotide metabolic process, oxidoreduction coenzyme metabolic process, L-proline biosynthetic process, glucose 6-phosphate metabolic process, endosperm development and others. In the MF analysis, responses to the defoliation mainly included glutamate 5-kinase activity, glucokinase activity, glutamate-5-semialdehyde dehydrogenase activity, unfolded protein binding, glucose-6-phosphate dehydrogenase activity, sucrose synthase activity, structural constituent of ribosome, amino acid kinase activity, pyruvate kinase activity, protein domain specific binding and chlorophyll binding. Defoliation mainly affected cytoplasm, endoplasmic reticulum lumen, intracellular part, cytoplasmic part, ribosome, chloroplast part and intracellular ribonucleoprotein complex (Fig. 5D; Table S2).

### Kyoto encyclopedia of genes and genomes (KEGG) Pathway Analysis

To understand the major responses of cellular processes to sink-source modifications in wheat flag leaves and grains, the KEGG pathway database was used with bases on the large-scale molecular dataset. In the pathway enrichment analysis, DEPs in half-degrained flag leaves were significantly enriched in linoleic acid metabolism and α-linolenic acid metabolism (in lipid metabolism), carbon fixation in photosynthetic organisms (in energy metabolism), nitrogen metabolism, carbon metabolism and vitamin B_6_ metabolism. Whereas, DEPs in flag leaves of defoliated plants were significantly enriched in photosynthesis, carbon metabolism, metabolic pathways, oxidative phosphorylation, glyoxylate and dicarboxylate and metabolism (Fig. 6A). In grains of half-degraining/intact group the DEPs were significantly enriched in four pathways in the KEGG database, carbon metabolism was the most represented pathway, followed by citrate cycle (TCA cycle), glycine, serine and threonine metabolism, glyoxylate and dicarboxylate metabolism and pyruvate metabolism, and most proteins in these pathways were decreased. Whereas, in grains of defoliation/intact group the DEPs were only enriched in two pathways including protein processing in endoplasmic reticulum and carbon metabolism in grains of defoliated plants (Fig. 6B). Taken together, the enrichment analyses based on the proteomics data suggested that photosynthesis, carbon metabolism, nitrogen/amino acid metabolism were the most sink/source manipulation responsive processes.

**Fig. 6.**
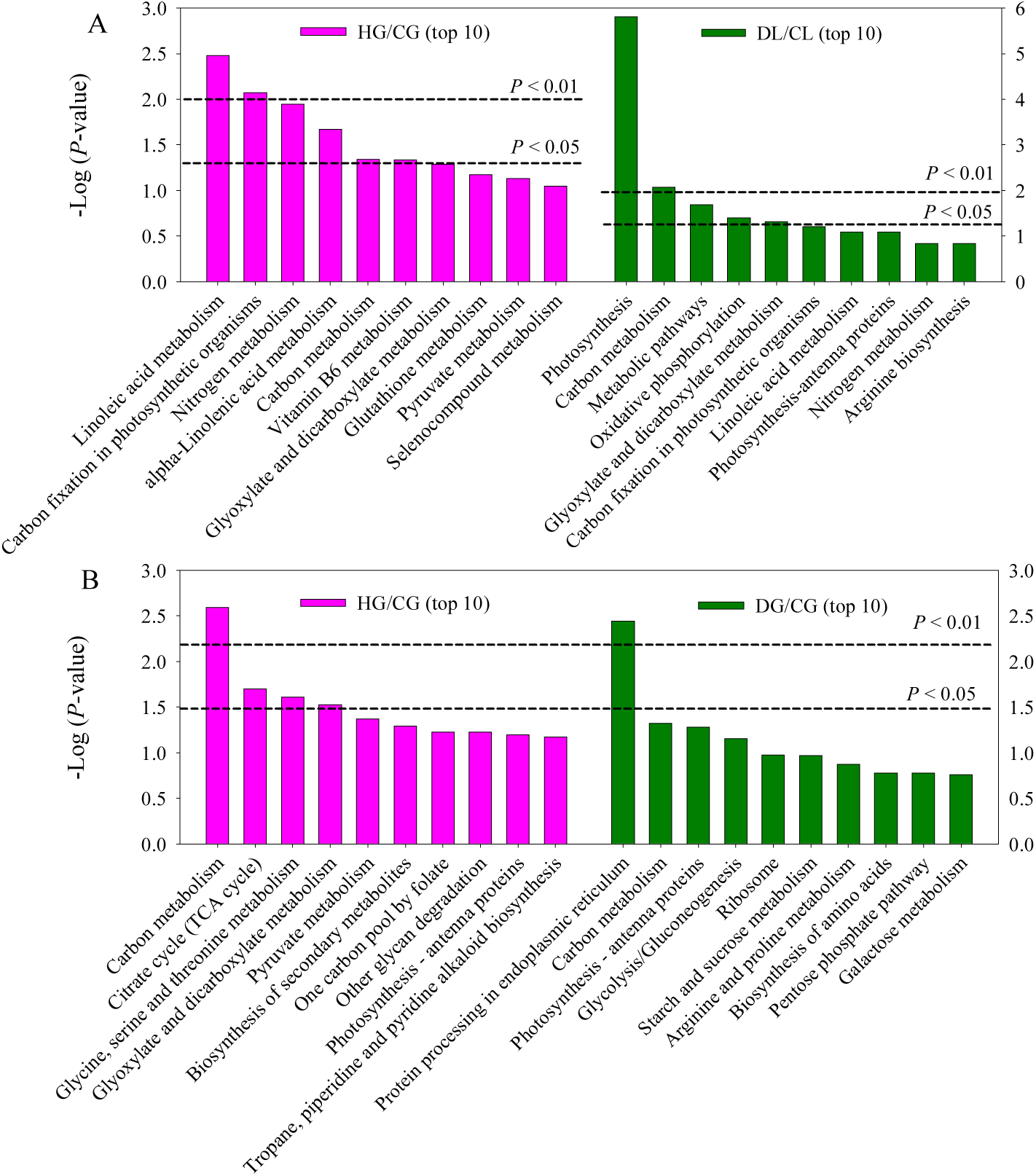
The most significant associated KEGG pathways of DEPs for HL/CL (flag leaf of half-degrained group compared with control) and DL/CL (defoliated group compared with control) (A) and HG/CG (grain of half-degrained group compared with control) and DG/CG (defoliated group compared with control) (B).

### PPI Analysis of DEPs

A regulation network of the DEPs identified in the current study was constructed to better understand the relationship between the proteins and affected pathways. As shown in Fig. 7, a complicated interaction network among DEPs was associated with the main biological processes.

**Fig. 7.**
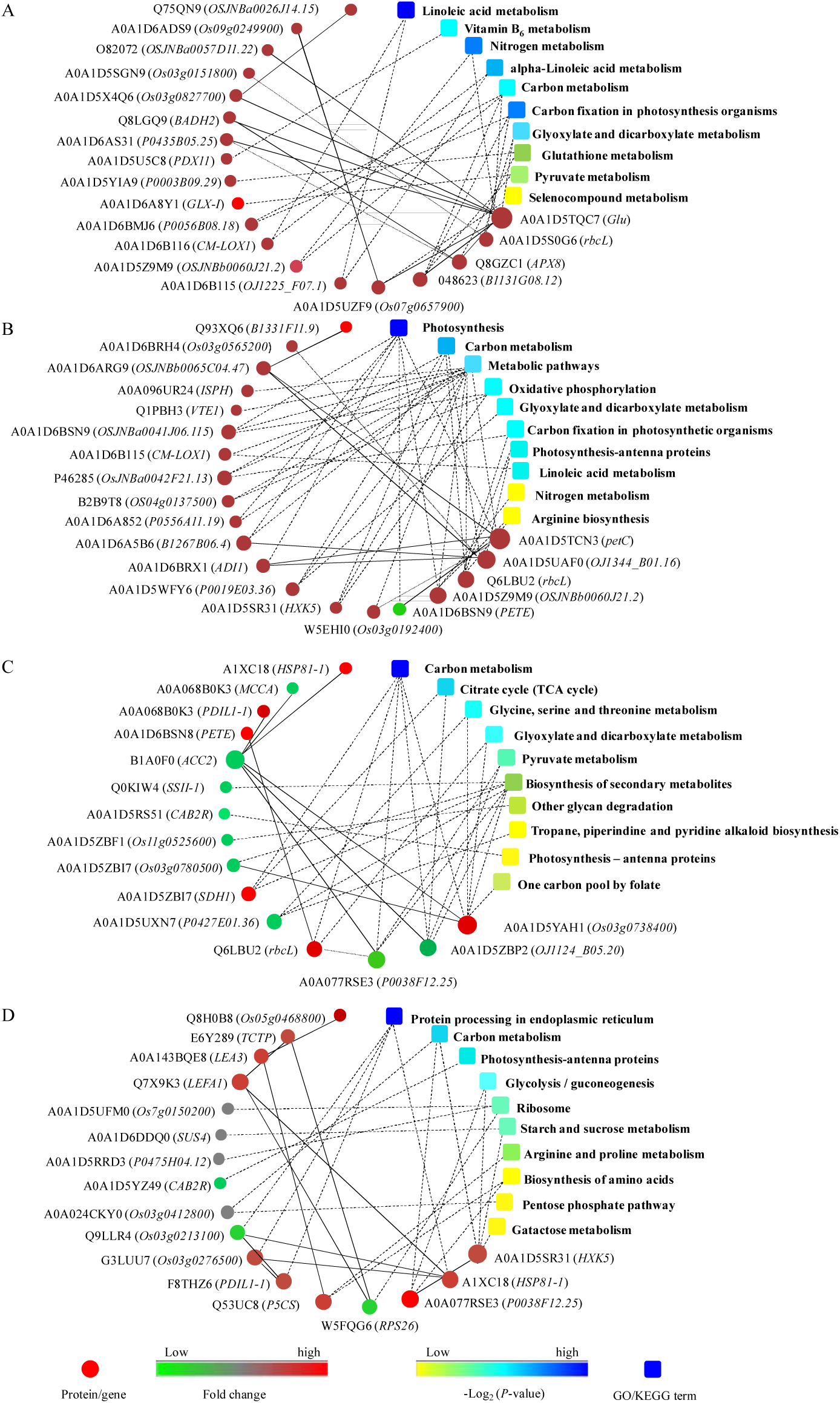
Interaction networks of the differentially expressed proteins in flag leaf of degrained wheat plants. Circle nodes denote differentially expressed proteins (genes) and colored rectangle indicates KEGG pathways. A solid line between two proteins indicates a known interaction annotated in the database; a dashed line between proteins indicates a potential interaction.

All the proteins constructed into the model for HL/CL were up-regulated. Main interactions were observed between A0A1D5TQC7 (*Glu*, encoding β-1,3-glucanase) and O82072 (*OSJNBa0057D11.22*, encoding phosphoenolpyruvate carboxylase), A0A1D5X4Q6 (*Os03g0827700*, encoding ATP-dependent RNA helicase), and Q8LGQ9 (*BADH2*, encoding betaine-aldehyde dehydrogenase), A0A1D6AS31 (*P0435B05.25*) and Q8GZC1 (*APX8*, encoding thylakoid-bound ascorbate peroxidase (fragment)), which were involved in the carbon metabolism, nitrogen metabolism, carbon fixation in photosynthesis organisms and glyoxylate and dicarboxylate metabolism; between Q8GZC1 and Q8LGQ9, and A0A1D5TQC7, and A0A1D6AS31, which were involved in the glutathione metabolism; between A0A1D5UZF9 (*Os07g0657900*, thioredoxin reductase) and A0A1D6ADS9 (*Os09g0249900*, encoding ferredoxin-thioredoxin reductase, catalytic chain), and A0A1D5TQC7, which were involved in the selenocompound metabolism. Many other interactions were found between proteins but remained characterized.

All the proteins constructed into the model for DL/CL were up-regulated except A0A1D6BSN9 (*PETE*). Main interactions were observed between A0A1D5UAF0 (*OJ1344_B01.16* encoding chlorophyll a-b binding protein) and A0A1D6ARG9 (*OSJNBb0065C04.47*, encoding peptidylprolyl isomerase), A0A1D6A5B6 (*B1267B06.4*, encoding putative oxygen-evolving enhancer protein 3-2, chloroplast (OEE3)), and A0A1D6BRX1 (*ADI1*, encoding ferredoxin) and A0A1D6BRH4 (*Os03g0565200*, HAD-superfamily hydrolase), which were involved in the carbon metabolism and photosynthesis - antenna proteins; between A0A1D5TCN3 (*petC*, encoding cytochrome b-c1 complex subunit Rieske) and A0A1D6ARG9, and A0A1D6BRX1, and W5EHI0 (*Os03g0192400*, encoding GRIM-19 family protein), and A0A1D6BSN9 (*PETE*, encoding plastocyanin), which were also involved in photosynthesis and carbon metabolism. Many other proteins were classified into biological functions including photosynthesis, oxidative phosphorylation, glyoxylate and dicarboxylate metabolism and nitrogen metabolism (Fig. 7B).

The DEPs for HG/CG were constructed into the model (Fig. 7C). Main interactions were observed between A0A1D5YAH1 (*Os03g0738400*, encoding serine hydroxymethyltransferase) and A0A1D6BSN8 (*(PETE*, encoding plastocyanin), B1A0F0 (*ACC2*, encoding plastid acetyl-CoA carboxylase (fragment)) and A0A1D5ZBI7 (*Os03g0780500*, encoding succinate dehydrogenase [ubiquinone] flavoprotein subunit), which were involved in five KEGG pathways including carbon metabolism, glycine, serine and threonine metabolism, glyoxylate and dicarboxylate metabolism, biosynthesis of secondary metabolites and one carbon pool by folate; between A0A1D5ZBP2 (*OJ1124_B05.20*, encoding dihydrolipoamide acetyltransferase component of pyruvate dehydrogenase complex) and B1A0F0, which were involved in carbon metabolism, citrate cycle, glyoxylate and dicarboxylate metabolism and biosynthesis of secondary metabolites; between A0A077RSE3 (*P0038F12.25*, encoding pyruvate kinase) and Q6LBU2 (*rbcL*, encoding rbcL gene product (30 AA) (fragment)), and B1A0F0, which was involved in three biological functions; between Q6LBU2 and A0A1D6BSN8, which was involved in two KEGG pathways. Many other proteins were classified into biological functions.

The DEPs for DG/CG were constructed into the model (Fig. 7D). Main interactions were observed between A1XC18 (*HSP81-1*, encoding heat shock protein XF20-1 (fragment)) and Q7X9K3 (*LEFA1*, encoding elongation factor-1 alpha (fragment)), and Q9LLR4 (*Os03g0213100*, encoding sec61p), and G3LUU7 (*Os03g0276500*, encoding cytHSP70 (fragment)), which were involved in protein processing in endoplasmic reticulum; between A0A1D5SR31 (*HXK5*, phosphotransferase) and A0A077RSE3 (*P0038F12.25*, encoding pyruvate kinase), which were involved in carbon metabolism, glycolysis/guconeogenesis, starch and sucrose metabolism and gatactose metabolism; among W5FQG6 (*RPS26*, encoding 40S ribosomal protein S26), E6Y289 (*TCTP*, translationally-controlled tumor protein) and Q7X9K3, which were involved in ribosome functions; between A0A143BQE8 (*LEA3*, encoding abscisic acid responsive protein Rab 15 (fragment)) and Q53UC8 (*P5CS*, encoding delta-1-pyrroline-5-carboxylate synthase), and Q8H0B8 (*Os05g0468800*, encoding cold regulated protein), which were involved in arginine and proline metabolism and biosynthesis of amino acids; between F8THZ6 (*PDIL1-1*, protein disulfide-isomerase) and Q9LLR4, and G3LUU7, which was involved in protein processing in endoplasmic reticulum. Many other proteins were classified into biological functions (Fig. 7D).

### Enzyme Activities

The activities of α-amylase and β-amylase in flag leaves were higher during 8 to 24 DAM than 0 DAM and remained constant during 8 to 24 DAM. Overall, α-amylase activity was lower than that of β-amylase. Compared with the intact plants, either half-degraining or defoliation did not affect the α-amylase activity at 8 and 16 DAM, but at 24 DAM defoliation significantly increased while half-degraining significantly decreased α-amylase activity. No difference in β-amylase activity was observed between half-degraining and control. However, defoliation significantly enhanced β-amylase activity at 16 and 24 DAM (Fig. 8).

**Fig. 8.**
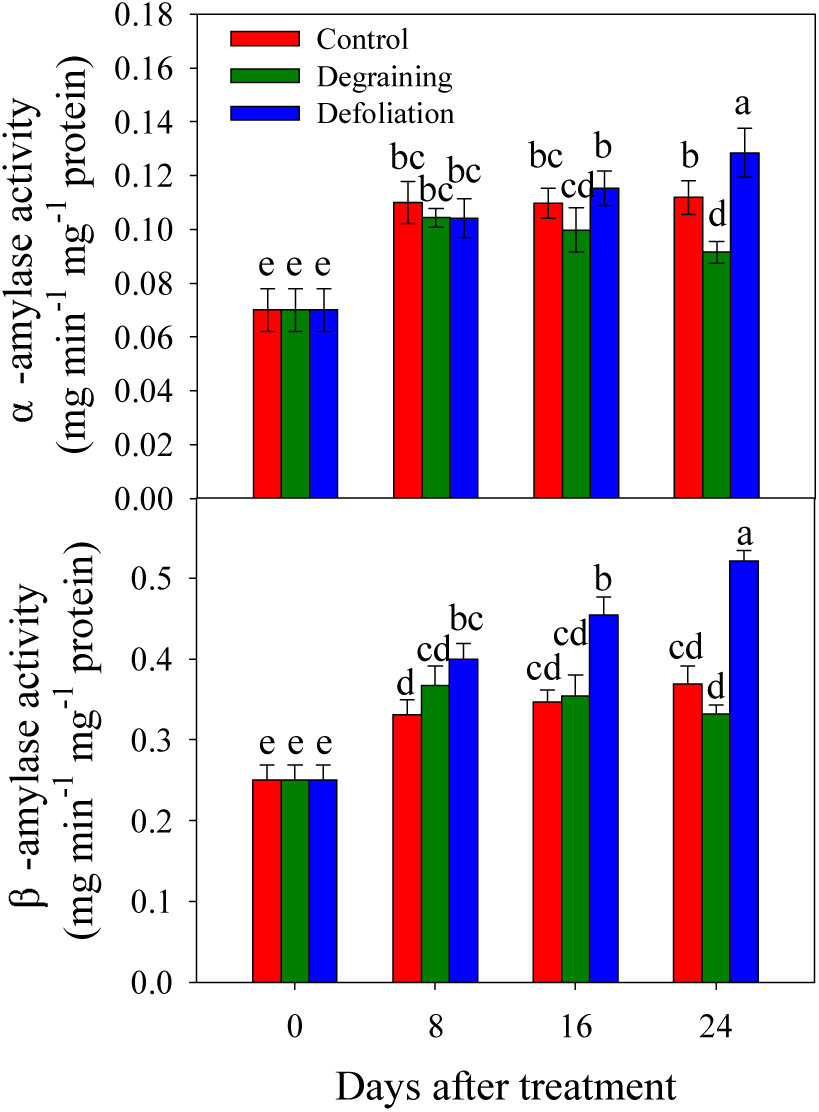
Comparison of the activities of α-amylase (a) and β-amylase (b) in flag leaf from the half-degraining, defoliation and control. Each value represents the mean ± SD from four independent samples. The columns labeled with different letters are significantly different at *p*<0.05 according to Duncan’s test for multiple comparisons.

During the experiment time, the activities of NP, AKP and ACP (Fig. 9) in flag leaves of intact plants remained relatively constant. Detraining decreased the NP, AKP and ACP activity at 8, 16 and 24 DAM compared with the intact plants. Defoliation decreased the NP activity at 8, 16 and 24 DAM but significantly increased the AKP activity at 8, 16 and 24 DAM and the ACP activity at 16 and 24 DAM.

**Fig. 9.**
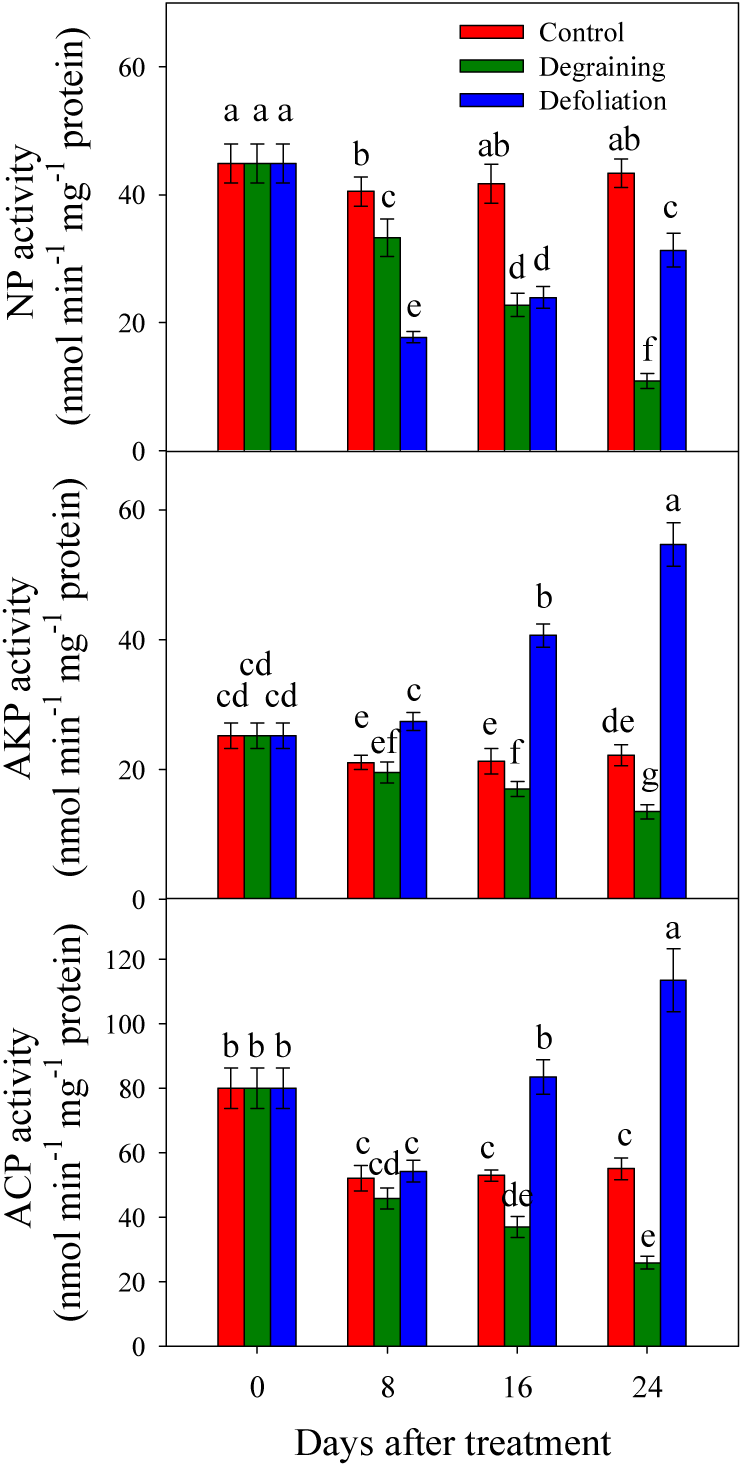
Comparison of the activities of neutral proteinase (a), acid proteinase (b) and alkaline proteinase (c) in flag leaf from the half-degraining, defoliation and control. Each value represents the mean ± SD from four independent samples. The columns labeled with different letters are significantly different at *p*<0.05 according to Duncan’s test for multiple comparisons.

### Grain Mass Responses to Half-degraining and Defoliation Manipulation

The removal of spikelets on one side of the spikes potentially doubled the resources available to the rest of the grains in the spikes; however, this manipulation did not halve the grain number per spike (Table 3). This de-graining increased the grain number per half-spike by 10.49% (*p*<0.05) and the single grain mass by 8.54% (*p*<0.05) compared with the in intact plants (control). As a consequence, half-degraining increased the grain mass per half-spike by 19.92%. Defoliation reduced the grain number per half-spike by 16.37% (*p*<0.05). The grain mass significantly responds to defoliation, causing reduction of the grain mass per half-spike by 19.64% (*P*<0.05).

**Table 3.** Effect of half-degraining and defoliation on the grain number per half-spike, single grain mass and number of spikelets and grain mass per half-spike.

|  | Grain number<br>half-spike<sup>-1</sup> | Single grain mass (mg) | Grain mass half-spike<sup>-1</sup> (mg) |
| --- | --- | --- | --- |
| Control | 17.35 b | 44.27 b | 768.08 b |
| Half-degraining | 19.17 a | 48.05 a | 921.12 a |
| Defoliation | 14.51 c | 42.54 b | 617.26 c |

## Discussion

The source/sink interaction modulates the leaf senescence and photosynthetic efficiency (Jessica *et al*., 2008) and thus determines the grain-filling rate and yield of cereals (Yang and Zhang, 2010). A strong source with sufficient reserves is necessary for grain filling and high sink capability promotes the reserve remobilization from source to sink (Fu *et al*., 2011). In the present study, sink-source manipulations were used to investigate the roles of source-sink relationship in regulation of leaf senescence and grain growth. Values of NDVI, PRI and SPAD and chlorophyll fluorescence were determined at a single-leaf basis and we observed that half-degraining increased but defoliation decreased the levels of these indices compared to the intact control (Table 1 and Fig. 1). Ultrastructural observations of flag leaf mesophyll cells at 16 DAM indicated that the mesophyll cells retained better for half-degraining (Fig. 2B) but inferior state for defoliation of the integrated chloroplasts (Fig. 2C). The NDVI, PRI and SPAD are the most frequently used spectral indices and are associated with chlorophyll contents and leaf senescence in higher plants (Chen and Chen, 2008). Chlorophyll fluorescence and chlorophyll degradation were also widely used as typical markers of developmental senescence progression and photosynthesis (Kong *et al*., 2010, 2017; Lyu *et al*., 2017; Kang *et al*., 2018). Higher values of them were closely associated with the grain yield (Vicente *et al*., 2018). The evidence from our physiological analysis and the ultrastructural observations of flag leaf mesophyll cells suggest that half-degraining may cause later but defoliation cause earlier leaf senescence.

It is well known that plant senescence is initiated and accelerated by active oxygen or free radicals. Antioxidant enzymes, such as SOD, POD and CAT, play important roles in scavenging excess ROS and reducing ROS detrimental effects on the cellular membrane. In wheat, the competence of the antioxidant defense system contributes the delayed senescence (Hui *et al*., 2012). The present study, the ROS content were higher in defoliated flag leaf than that of intact plants, although the defoliation significantly increased the activities of SOD, POD and CAT in the flag leaves, especially at 24 DAM. In DEPs identified using iTRAQ analysis, SOD (Fragment) was upregulated in defoliated plants at 16 DAM (Table S1). These results indicate that partial source removal may induce more active physiological responses and thus cause a burst of ROS and membrane oxidation which in turn result in the premature senility of the wheat plants. The increased activity of these antioxidant enzymes may be a strategy to protect plant from more serious damage, partially relieve the plant senescence and attempting to increase the active grain-filling period and the grain mass. On contrary, in half-degrained flag leaves, the activities of SOD, POD and CAT decreased but the ROS content also slightly decreased (Fig. 3; Table S1 for POD), indicating that half-degraining lay a lower “burden” to flag leaf and lower ROS production.

Plant hormones and associated hormonal crosstalk are significantly related to the modification of source/sink relationships, a stronger sink capacity, grain filling rate (Yang *et al*., 2001; Yang and Zhang, 2006; Liu *et al*., 2016), membrane stability, foliar senescence and photosynthetic efficiency, possibly by acting as signalings to scavenge ROS or prevent ROS formation (Yang *et al*., 2002; Afzal *et al*., 2005; Biswas and Mondal, 1986).

Cytokinins are the most significant phytohormones regulating the initiation and timing of leaf senescence (Xu and Huang, 2007; Reguera *et al*., 2013; Jan *et al*., 2019) and closely associated with the progress of leaf senescence in the stay-green mutant *tasg1* of wheat (Wang *et al*., 2016) and in the stay-green phenotype of maize (*Zea mays* L.) (He *et al*., 2005). The grain CTK concentrations were significantly and positively correlated with the grain-filling rate and maximum grain mass (Yang *et al*., 2002; Yang *et al*., 2008; Zhang *et al*., 2009). In rice (*Oryza sativa* L.), wheat, maize and barley (*Hordeum vulgare* L.), a higher CTK concentration was generally observed in the endosperm of the grains and involved in cell division during the early phase of grain development which, in combination with functions of auxin, are positively and significantly correlated with cell division, sink strength and the grain-filling rate (Dietrich *et al*., 1995; Yang *et al*., 2000; Zhang *et al*., 2016; Xu *et al*., 2007; Zhang *et al*., 2009a; Liu *et al*., 2016; Yang *et al*., 2016), due to the enhanced sink strength (Rijavec *et al*., 2009; Yu *et al*., 2015).

High IAA concentrations in the grains could create an “attractive power”, leading to an increase in CTK concentrations in grains (Seth and Waering, 1967; Singh and Gerung, 1982). Lur and Setter (1993) reported that grain IAA content rapidly increased in maize at nearly grain filling, which was associated with an increase in assimilate transport to the developing grains. Auxin is linked with enhanced sink capacity, resulting in higher grain-filling rate, enlargement of cell, increased nutrient assimilation and source nutrient remobilization (Kong *et al*., 2015). In cucumber, leaf auxin promotes source N remobilization during the reproductive stage (Du *et al*., 2018).

GA signaling implicated for regulatiing leaf senescence; knockout of the GA biosynthesis gene *GA REQUIRING1* (*GA1*) results in delayed onset of senescence, because GA activates α-amylase gene (Chen *et al*., 2014). While in grain, enhanced levels of GA_3_ or in combination with CTKs, were found to be beneficial for enhanced grain filling and grain size (Wang *et al*., 2006; Suzuki *et al*., 2012).

In the present study, we observed that defoliation decreased the contents of CTKs and IAA in flag leaf (except at 8 DAM); however, this manipulation significantly increased the contents of CTKs, IAA and GA_3_ in grain and thus increased the grain/leaf ratio of CTKs, IAA and GA_3_ contents. Based on the results from current and previous studies, we postulate that higher source completion due to defoliation stimulate the hormone biosynthesis, thereby trying to increase sink strength and enhancing their potential to absorb more of the carbohydrate transported from the limited source. Meanwhile, decreased levels of CTKs and IAA for defoliation promote the degradation of large molecules for grain filling, which cause early senescence as demonstrated by ultrastructural observation and measurement of vegetation indices (Table 1; Fig. 2). This view is highly consistent with the findings that in cultivar with relative large sink ratio exists a greater competition for source photosynthate supply to meet the grain filling, leading to premature senescence of the plant (Yang and Zhang, 2010). Furthermore, the data presented here suggest that that the ratio of grain to leaf CTK, IAA and GA_3_ contents may be the decisive factor in regulating leaf senescence in wheat.

Half-degrainning also significantly decreased the CTKs content in flag leaf at 8 and 16 DAM and in grain at three time points, decreased the IAA content at 16 DAM in flag leaf but show no effect on that in grain, and decreased GA_3_ content in flag leaf but significantly increased the GA_3_ content in grain (Table 1; Fig. 2). Further analysis indicated that half-degraining decreased the grain/leaf ratio of CTK content. Based on these results, we suggest that half-degraining may result in the lower sink strength and higher contents of CTKs and IAA may not be required to compete for source reserves and therefore resulting delayed leaf senescence. Contrary to the data for defoliation, the lower ratio of grain to leaf CTK content may contribute the delayed leaf senescence for half-degraining. As for GA_3_, the lower content was consistent with the decreased α-amylase activity in flag leaf and higher GA_3_ stored in the grain may be favorable for its germination.

SA, JA, CTKs and other hormones are the major hormones that have antagonistic and synergistic signaling effects on plant senescence (Thomas and Ougham 2014). Accumulation of ROS and JA lead to early leaf senescence and low photosynthesis efficiency (Wang *et al*., 2015). Specific inhibition of JA biosynthesis in the leaf resulted in grain yield increase in rice (Tamaki *et al*., 2015). The connection between leaf senescence and SA is still unclear. However, increasing body of evidence has demonstrated that SA in wheat act as a signaling molecule to induce an endogenous ROS detoxification, increases the activities of SOD, CAT, APX and glutathione reductase, help preserve photosynthetic function in stresses (see review by Abhinandan *et al*., 2018). In our study, we observed that in flag leaf, defoliation increased the free SA but decreased the JA content during experimental period. While in grain, exactly reverse effects of both manipulations on SA and JA levels were observed (Table 2). The current results indicate that antagonistic action between JA and SA may exist in regulating leaf senescence and grain filling, and the significantly higher SA/JA ratio (data not shown) may help to protect plant from too early leaf senescence of defoliated plants. Half-degraining increased the SA content at 8 and 16 DAM but decreased the JA content at 8 and 16 DAM in flag leaf, increased the SA content at 8 DAM but decreased the JA content at 8 and 16 DAM in grain. We hypothesized that the SA/JA interplay might be is possible to function in delaying the initiation of leaf senescence and promote the early grain setting in half-degrained plants.

Leaf senescence is associated with fundamental changes in the proteome. In the present study, a comparative proteome analysis based on iTRAQ was performed to identify the critical candidate factors involved in leaf senescence and grain filling. GO terms within the biological process category, the molecular function category and cellular components were showed in Fig. 5. KEGG pathway enrichment analysis of these genes and PPI were also performed (Fig. 6).

The destruction of the oxidation-reduction system is accompanied by an increase in burst of ROS such as superoxide (O^2−^) and hydrogen peroxide (H_2_O_2_), which may lead to early leaf senescence (Li *et al*., 2014; Parihar *et al*., 2015). Glutathione metabolism can reduce and eliminate oxidative damage caused by ROS plays an important role in maintaining redox balance (Hicks *et al*., 2007). In this pathway, glutathione is recycled through the oxidation/reduction process and thus involved in oxidative stress. Ferredoxin:thioredoxin reductase can acts as one component of antioxidative defence systems (Geigenberger *et al*., 2017) and was required for proper chloroplast development and was involved in the regulation of plastid gene expression in *Arabidopsis thaliana* (Wang *et al*., 2014). In DEPs of our study, ferredoxin:thioredoxin reductase was upregulated in the half-degraining leaf and was clustered into the MF. The chloroplast is the central organelle that produces ROS, whereas accumulation of ROS may cause oxidative damage and inhibit photosynthesis (Exposito-Rodriguez *et al*., 2017). In comparison between HL and CL group, the most prevalent GO terms for BP were involved in the ROS metabolic processes, such as oxidation-reduction process, H_2_O_2_ catabolic process, response to oxidative stress, H_2_O_2_ metabolic process, O^2−^ removal and so on (Table S1). In these processes, the homeostasis of oxidation-reduction process was most strongly enhanced in degrained flag leaf as judged by the PAS_Zscore and oxidoreductase activity (MF) and activities of several kinds of oxidoreductase (MF) were significantly affected, which may occur in the chloroplasts as indicated in the CC analysis (Fig. 5A) and glutathione metabolism was constructed to the top 10 of KEGG pathways for HL/CL. In MF analysis for HL/CL, phytoene dehydrogenase acitivty was significantly affected, which was associated with carotenoid biosynthetic pathway, serve as membrane integrated antioxidants, protecting cells from oxidative stress (Liu *et al*., 2012). The thylakoid-bound ascorbate peroxidase (Fragment) and betaine aldehyde dehydrogenase were important factors in abiotic stresses and can also function as a ROS scavenger (Golestan Hashemi *et al*., 2018) and both were involved in many pathways as indicated in PPI analysis (Fig. 7) and upregualted in flag leaf of half-degrained plants (Table S1). These results imply that half-degraining may enhance antioxidant ability by promoting ROS scavenging activities and at least partially contributed to higher chlorophyll content (as indicated by higher SPAD values) and delayed leaf senescence.

KEGG pathway analysis for HL/CL suggested that the DEPs were mainly involved in linoleic acid metabolism (lipoxygenase, up-regulated), nitrogen metabolism (such as upregualted biological process organonitrogen compound metabolic process and organonitrogen compound biosynthetic process). Correspondingly, MFs including glutamate synthase activity, glutamate synthase (ferredoxin) activity and pyridoxal 5’-phosphate synthase (glutamine hydrolysing) activity were affected and carbon metabolism (carbohydrate biosynthetic process, starch biosynthetic process, starch metabolic process, cellular carbohydrate biosynthetic process, carbohydrate metabolic process, carboxylic acid metabolic process. The MFs such as phosphoenolpyruvate carboxylase activity, phosphoenolpyruvate carboxykinase activity, sucrose-phosphate phosphatase activity were affected (Table S1). Of the most interesting, sucrose phosphate phosphatase was associated with source activity, enhancing transport of its resulting product sucrose from the photosynthetic tissues via the phloem into sink (Maloney *et al*., 2015) and thus affecting progress of leaf senescence in wheat (Wang *et al*., 2016). BP analysis showed that these processes were upregulated, indicating that in half-degrained flag leaf, nitrogen and carbon metabolism may be enhanced.

As for the cellular component, categories playing important roles in protein biosynthesis such as the cytosolic ribosome and ribosome were highly represented in DL (Fig. 5C; Table S2). Accordingly, DEPs such as 60S ribosomal and 40 S ribosomal proteins were upregualted (Table S1).

At the same time, DEPs related with photosynthesis (carbon fixation) and generation of precursor metabolites and energy that occur in the thylakoid, plastid) were upregualted (Table S1). For example, phosphoenolpyruvate carboxylase was upregulated, which showed an significant interaction with carbon fixation in photosynthesis organisms (Fig. 7).

In the grains of half-degrained plants, amine oxidase (DEPs, A0A1D5UXN7, A0A1D5UXN8, A0A1D5UXN9, A0A1D6DCS6), which catalyzes the reaction to produce H_2_O_2_, was down-regulated and the MF of primary amine oxidase activity (GO:0008131) was affected. Superoxide dismutase [Cu-Zn] (A0A1D5TQL0, A0A1D5TQL1, A0A1D5TQL3, W5B1E5) and dehydroascorbate reductase was upregulated. Oxidoreductase activities (GO:0016635, GO:0016721, GO:0016641, GO:0016864, GO:0016638) were also affected. Protein glutathionylation (GO:0010731), a mechanism for redox regulation and signaling (Gallogly and Mieyal, 2007), was upregulated in the current study. These results indicate the decrease in oxidative stress to degrained plants.

The significantly affected KEGG pathways mainly included the carbon and nitrogen metabolism. However, nearly all the nitrogen- and carbon-related in the top 20 of BP were inhibited (Table S2). Most of biological processes related to metabolites and energy were significantly downregualted. This response indicates that low levels of carbohydrate (derivatives) are needed to be processed in the grains of half-degrained plants.

Protein disulfide isomerases (PDIs) are enzymes found primarily in the endoplasmic reticulum (ER) in eukaryotes play a vital role in protein folding. The up-regulated levels of PDIs (A0A024FRN3, A0A1D5XQP7, A0A1D5XQP9, B9A8E3, F8THZ6, Q6JAB5, Q93XQ8) in the present study indicated the higher PDI activity (MF) and confirmed the active status of protein synthesis (Hayano *et al*., 1995; Lv *et al*., 2016; Kayum *et al*., 2017) and higher resistance to abiotic stresses (Kayum *et al*., 2017). This conclusion was strongly confirmed by the upregulation of 60S ribosomal protein L29 (A0A1D5RRD3), 40S ribosomal protein S12 (A0A1D5UFM0, A0A1D5UVS7) and Eukaryotic translation initiation factor 4B1 (Q9AUJ7) (Table S1).

In plants, acetyl-CoA carboxylase activity is found in the plastids and catalyzes the first committed step of de novo fatty acid biosynthesis by the carboxylation of acetyl-CoA to malonyl-CoA (Fazli *et al*., 2005; Liu *et al*., 2007; Lv *et al*., 2016). In the present study, acetyl-CoA carboxylase (Q8L4W1), plastid acetyl-CoA carboxylase (fragment) (A0A068B0K3, A0A068B0L8, A0A068B0M7, A0A068B226, A0A068B2H1, A0A068B2I1, A0A068B3I5, A0A068B675, B1A0F0, D0V7D7, D0V7D9, D0V7E1, D0V7E3, D0V7F1) and acetyl-coenzyme A carboxylase (Fragment) (O48960), were down-regulated, indicating that CoA carboxylase activity (GO:0016421), acetyl-CoA carboxylase activity (GO:0003989) and biotin carboxylase activity (GO:0004075) were affected. In PPI analysis, B1A0F0 (ACC2) showed interactions with many other DEPs, indicating fatty acid biosynthesis may be downregulated and many pathways were affected (Fig. 7).

Processes involved in photosynthesis such as photosynthesis (GO:0015979), photosynthesis and light reaction (GO:0019684), photosynthesis, light harvesting (GO:0009765) and photosynthesis, dark reaction (GO:0019685) were also significantly up-regulated in the grain of degrained plants, indicating that higher use efficiency of light, CO_2_ and supplication of higher amount of O_2_ to the developing endosperm (Kong *et al*., 2016). In addition, interaction of plastocyanin (A0A1D6BSN8) with rbcL gene product (30 AA) (fragment) (Q6LBU2) was observed and both DEPs were upregulated, indicating that carbon metabolism and glyoxylate and dicarboxylate and metabolism were enhanced (Fig. 7).

In the molecular function category, processes was involved in succinate dehydrogenase activity (GO:0000104) and succinate dehydrogenase (ubiquinone) activity (GO:0008177) (Fig. 5C). These results were in accordance with upregulation of succinate dehydrogenase [ubiquinone] flavoprotein subunit, mitochondrial (A0A1D5ZBI7) and represented MF category such as mitochondrial electron transport, succinate to ubiquinone (GO:0006121) (Fig. 5C) and citrate cycle (TCA cycle) (Fig. 6B). These processes occurred in mitochondrial matrix (GO:0005759), mitochondrial respiratory chain complex II (GO:0005749), succinate dehydrogenase complex (ubiquinone) (GO:0045257) (Fig. 5C). In addition, GMP biosynthetic process (GO:0006177) and GMP metabolic process (GO:0046037) were significantly represented. In the molecular function category, processes involved in hydrolase activity (hydrolyzing O-glycosyl compounds) (GO:1901657), glycosyl compound metabolic process (GO:1901659) and glycosyl compound biosynthetic process (GO:1901659) were significantly represented (Table S2). These results indicate that the energy generation may be increased in grain of degrained plants.

In our study, relative ROS content increased in defoliated flag leaf (Fig. 3), which may inhibit photosynthesis (Exposito-Rodriguez *et al*., 2017) as discussed above. However, in defoliation plants glutathione s-transferase (W5E8I6), superoxide dismutase (Fragment) (A3FKE5), oxidoreductase activity, acting on superoxide radicals as acceptor (GO:0016721), superoxide dismutase activity (GO:0004784) (Fig. 3), oxidation-reduction process (GO:0055114), oxidoreduction coenzyme metabolic process (GO:0006733) (Table S2) the tocopherol cyclase (Q1PBH3) and glutathione S-transferase (W5E8I6) and thus tocopherol cyclase activity (GO:0009976) were upregulated, indicating that in the flag leaf of defoliated plants, some strategies were developed to eliminate oxidative damage caused by ROS (Kanwischer *et al*., 2005; Liang *et al*., 2018). However, these means can only partially compensate for the ROS burst, because the overall capability of ROS metabolic process (GO:0072593), removal of superoxide radicals (GO:0019430), cellular response to superoxide (GO:0071451, GO:0000303), cellular response to oxygen radical (GO:0071450, GO:0000305), regulation of H_2_O_2_ metabolic process (GO:0010310) and cellular response to ROS (GO:0034614) were downregulated (Table S2).

Interestingly, the top 50 significantly represented BPs were upregulated in flag leaf of defoliated plants. GO terms relative to the photosynthesis were significantly enriched, but showed different changing trends. Plastid organization (GO:0009657) and cofactor metabolic process (GO:0051186) were upregulated. In the PPI net works, DEPs related to photosynthesis-antenna proteins and carbon metabolism were also upregulated (Fig. 7). However, stomatal complex morphogenesis (GO:0010103), stomatal complex development (GO:0010374), chlorophyll biosynthetic process (GO:0015995) chlorophyll metabolic process (GO:0015994), chlorophyll biosynthetic process (GO:0015995) and pigment biosynthetic process (GO:0046148) were downregulated (Table S2). As a consequence, the net photosynthetic efficiency decreased (Fig. 5B). These changes occurred in chloroplast (CC). In MF analysis, iron-sulfur cluster binding, 2Fe-2S iron-sulfur cluster binding and 3Fe-4S iron-sulfur cluster binding play an important role in photosynthetic rate by regulating electron transfer and chlorophyll content in rice (Zhao *et al*., 2015), however, these GO terms were significantly affected and thereby influencing electron carrier activity, proton-transporting ATPase activity, rotational mechanism, hydrogen-exporting ATPase activity, hydrogen ion transmembrane transporter activity, ATPase activity, plastoquinol-plastocyanin reductase activity (located in plastid and chloroplast thylakoid membrane). These functions were closely coupled to transmembrane movement of ions, rotational mechanism, monovalent inorganic cation transmembrane transporter activity, electron transporter, transferring electrons from cytochrome *b6/f* complex of photosystem II activity (Fig. 5B). The plastoquinol-plastocyanin reductase activity (GO:0009496) and electron carrier activity and electron transporter, transferring electrons from cytochrome b6/f complex of photosystem II activity (GO:0046028) that are located in the chloroplast cytochrome ***b6-f*** complex (GO:0009512) enables the directed movement of electrons from the cytochrome *b6/f* complex of photosystem II and couples the energy released from electron transport to proton translocation across the membrane. Therefore, these results indicate that in the flag leaf of defoliated plants, energy supply may be very important. Indeed, The GO categories enriched are primarily related to energy metabolism were upregulated, including glyceraldehyde-3-phosphate metabolic process (GO:0019682), glucose 6-phosphate metabolic process (GO:0051156) and carbohydrate derivative metabolic process (GO:1901135) and generation of precursor metabolites and energy, pentose-phosphate pathway, pentose-phosphate shunt).

The accumulation of peptidyl-prolyl cis-trans isomerase (Q93XQ6), peptidylprolyl isomerase (A0A1D6ARG9) mediates protein folding (GO:0006457) and interacts with chlorophyll a-b binding protein and cytochrome b-c1 complex subunit Rieske (Fig. 7), thus promoting the functions of these proteins. As for lipid metabolism, organophosphate metabolic process (GO:0019637), cellular lipid metabolic process (GO:0044255, GO:0006629), lipid biosynthetic process (GO:0008610) and phospholipid biosynthetic process (GO:0008654, GO:0006644) were also upregulated.

In defoliated grain, protein glutathionylation (GO:0010731), a mechanism for redox regulation and signaling (Gallogly and Mieyal, 2007), and oxidation-reduction process (GO:0055114) were downregulated in the current study, indicating a decrease in the ability to scavenge ROS.

In the BP category analysis, organonitrogen compound metabolic process (GO:1901564) and cellular protein metabolic process (GO:0044267) were enhanced, which may be consistent with the increases in proteinase activities (Fig. 9). However, protein folding in endoplasmic reticulum (GO:0034975), protein folding (GO:0006457) were downregulated (Fig. 5); accordingly, protein processing in endoplasmic reticulum was significantly affected (Fig. 6) was downregulated. sec61p (Q9LLR4) In further, translation (GO:0006412), peptide biosynthetic process (GO:0043043), peptide metabolic process (GO:0006518), and L-proline biosynthetic process (GO:0055129, GO:0006561) were downregulated. Accordingly, a large number of low molecular weight glutenin (subunits) were decreased (Table S2). These processes may occur in endoplasmic reticulum lumen (GO:0005788), prefoldin complex (GO:0016272) and ribosome (GO:0005840), regulating structural constituent of ribosome (GO:0003735) (Table S2).

Glucose 6-phosphate metabolic process (GO:0051156) and generation of precursor metabolites and energy (GO:0006091) were downregulated. Meanwhile, pyruvate kinase (A0A077RSE3, A0A1D5WW05, A0A1D5WW06, A0A1D6RQH6) (Table S1) and thus pyruvate kinase activity (GO:0004743) decreased, suggesting that energy generation was decreased in the grain of defoliated plants.

Chlorophyll a-b binding protein, chloroplastic (A0A1D5YZ49, A0A1D5YZ50, A0A1D5ZLT0, A0A1D6S1V3) was downregulated. Moreover, photosynthesis, light harvesting (GO:0009765) and photosynthesis, light reaction (GO:0019684) were also decreased. These processes might occur in the chloroplast parts (GO:0009707, GO:0044434, GO:0009941, GO:0031969). These results were consistent with the decrease in photosynthetic efficiency (Fig. 1).

From these results, we surmised that defoliation might have aggravated the premature senescence of the wheat plants and ultimately, endosperm development (GO:0009960, GO:0048316), seed development (GO:0048316) and seed maturation (GO:0010431, GO:0010431) were inhibited.

A large amount of macromolecules in normal senescent leaves are degraded into small molecules by degradation enzymes (Buchanan-Wollaston *et al*., 2003). The translocation of non-structural carbohydrates in leaf to grain are essential for growing grains, which was closely correlates with senescence (Smith *et al*., 2005). Starch is the main carbohydrates and primarily hydrolyzed by a- and β-amylase. Genes encoding proteases to hydrolyze proteins show highly induced expression during leaf senescence in different species (Liu *et al*., 2008; Desclos *et al*., 2009; Hollmann *et al*., 2014; Moschen *et al*., 2016). At least a portion of senescence-associated proteases localize to senescence-associated vacuoles to degrade chloroplast-derived proteins and are up-regulated during leaf senescence and then the resulting catabolic products are mobilized from leaves to the developing seeds (Drake *et al*., 1996; Buchanan-Wollaston *et al*., 2003; Carrión *et al*., 2013; Li *et al*., 2017). In the present study, defoliation increased the activities of a-amylase, β-amylase (Fig. 8), acid proteinases and alkaline proteinases (Fig. 9) but decreased the activity of neutral proteinases. Half-degraining decreased the activities of proteinases while not affected the activities of a-amylase and β-amylase. Nevertheless, these results indicate that the earlier senescence of defoliated plants might be due to the increases in these hydrolytic enzymes, while the delayed leaf senescence of half-degrained plants might be caused by the decreased starch degradation.

Vegetative organ CTKs play an important role in regulating senescence which was associated with a delay of proteolytic activity and thus the N remobilization to developing grains (Noodén *et al*., 1997; Roberts *et al*., 2011; Gregersen *et al*., 2013). Therefore, we speculate that the decrease in the CTK content may be associated with the increases in proteolytic activity in flag leaf of defoliated plants.

In conclusion, reduced sink/source ratio due to half-degraining delayed wheat leaf senescence while higher sink/source ratio due to defoliation caused higher amount of ROS production and facilitated the degradation of chlorophyll proteins, carbohydrate and thus the plant senescence. CTKs, IAA, GA_3_, JA and SA and interactions among these hormones had major roles in regulating source capability and sink strength and thereby impacting the process of leaf senescence. Sink and source manipulations induces a lot of deferentially expressed proteins, which were mainly involved in ROS scavenging, leaf photosynthesis, carbon and nitrogen metabolism, generation of precursor metabolites and energy and grain development. Determination of hydrolytic enzyme activities further suggested that degradation of macromolecule compounds of carbon and nitrogen was promoted in flag leaf of the defoliated plants but retarded in the half-degrained plants, which in turn impacted the carbon and nitrogen metabolisms in the grain. Although trimming the spikes did not produce significant changes in duration of grain filling, the single grain growth was enhanced by half-spike removal and depressed by defoliation, indicating that the yield potential of wheat is limited by both sink capacity and source availability. Our results indicate that future yield improvements may be achieved by strengthening both the source and sink capacity in breeding for increased yield potential in wheat.

## Material and Methods

### Site, experiment and design

The experiments were conducted in 2017-2018 in a field at an experimental station (36°42′N, 117°4′E; altitude 48 m) of the Shandong Academy of Agricultural Sciences, China. The soil was a fine loamy. Wheat cultivars Jimai 23 was sown at a seeding rate of 10 g m^−2^ on October 8. Plots were fertilized before planting with 10 g m^−2^ P_2_O_5_, 10 g K_2_O m^−2^ and 7.5 g N m^−2^. At the shooting stage (Zadoks stage 31), 15 g N m^−2^ was top-dressed with urea.

### Source and Sink Manipulations

To analyze the effects of sink and source reduction on grain growth, sink-source manipulations were conducted as follow: (1) half of the spikelets were removed from one side of the spikes by hand to double the assimilate availability for the remaining grains; (2) all leaves of culms except flag leaf were removed to reduce the source assimilate availability. Each manipulation was performed in three 1-m sections of the rows at 2 days after anthesis. Three 1-m sections were selected and were left intact as a control.

### Measurement of Photochemical Reflectance Index (PRI), Normalized Difference Vegetation Index (NDVI) and the Soil-plant Analyses Development (SPDA) Values of Flag Leaves

Leaf NDVI and PRI values were measured with a PlantPen instrument (Photon Systems Instruments, Brno, Czech Republic) on 30 leaves per plot. The SPAD index were determined on the at least 30 flag leaves per plot using a chlorophyll meter (SPAD-502 plus, Konica Minolta, INC. Japan) at 8, 16 and 24 days after manipulation (DAM). Data from each plot were averaged to obtain a mean.

### Chlorophyll Fluorescence Assay and Imaging

Chlorophyll fluorescence analysis was performed at different stages to determine the maximum PSII quantum yield (*Fv*/*Fm*) and the effective PSII quantum yield (Φ_PSII_) in the flag leaves using a kinetic imaging fluorometer (FluorCam, Photon System Instruments Ltd., Brno, Czech Republic) as described by Kong *et al*. (2000).

### Transmission Electron Microscopy

Five flag leaves were randomly collected from three plots of each treatment and the middle part of the leaf blade was sectioned into 2 m × 2 m patches using a razor blade. The collected samples were immediately fixed in 2.5% glutaraldehyde solution in 100 mM phosphate buffer (PB, pH 7.2) and stored for 14 h at 4°C. After washing with the buffer, the samples were post-fixed with 1% (w/v) OsO4 in the same buffer at 4°C for 4 h. The samples were then dehydrated in an ethanol series, transferred into propylene oxide and finally embedded in polymerized Epon812 resin. Ultrathin sections were cut with an LKB-V microtome and then mounted on a formvar-coated brass grid. The sections were double stained in 2% uranyl acetate (w/v) in 70% methanol (v/v) and the in 0.5% lead citrate. The ultrastructures were observed and imaged using transmission electron microscopy (JEM-1200EX; JEOL Ltd., Tokyo, Japan) at 80 kV.

### Hormone Analysis

Flag leaves or spikes in triplicate of each treatment were collected at 8, 16 and 24 DAM, immediately frozen in liquid nitrogen and then stored at −80°C and then the leaves and grains were used for each hormone analysis. The hormone was extracted using extraction kit and the contents of phytohormones were measured following the instructions of manufacturer (SuZhou Comin Biotechnology; Suzhou, China).

Briefly, for the extraction and purification of zeatin, zeatin ribuside, kinetin, GA_3_ and IAA, sample (approximately 0.10 g flag leaves or grains) was ground in a mortar (on ice), and 5 ml 80% (v/v) methanol extraction solution containing 1 mM butylated hydroxytoluene was used as an antioxidant. The methanolic extracts were incubated at 4°C over night and centrifuged at 10,000 g for 15 min at 4°C. The supernatants were dried with N_2_ at 40°C, dissolved in 200 μL methanol and filtered through a 0.45-μm membrane. High Performance Liquid Chromatography (HPLC; Rigol L3000, Beijing, China) with Kromasil C18 reverse phase column was used to measure the hormone content. The mobile phase was prepared by mixing methanol and ultrapure water at a ratio of 2:3 (v/v). Injection volume was 10 μL, flow rate 0.8 ml/min, column temperature 35 °C, aliasing time 60 min, and detection wavelength 254 nm. Three independent biological replicates of each sample were performed.

For SA and JA analysis, sample (approximately 0.10 g) was grounded to fine powder in liquid nitrogen and extracted with 1.0 ml of 90% methanol over night at 4°C. After centrifugation at 8,000 × g for 10 min, the precipitate was re-extracted with 0.5 ml of 90% methanol for 2 h and re-centrifuged. The supernatants from both extractions were combined and air-dried in a water bath at 40°C. The dried samples were resuspended in solvent containing 20 μL of 1 mg/ml trichloroacetic acid and 1 ml of ethylacetate/cyclopentane (1:1, v:v) by vigorous vortexing for 30 min and centrifuged at 8,000 × g for 10 min. The top organic phase (including SA and JA) was dried, dissolved in mobile phase and injected into a reverse phase C18 HPLC column equipped with a fluorescence detector for the analysis of SA measured. For SA, the mobile phase was prepared by mixing methanol and ultrapure water at a ratio of 2:3 (v/v). Injection volume was 10 μL, flow rate 0.8 ml/min, column temperature 35°C, aliasing time 45 min and monitored with excitation at 294 nm and emission at 426 nm. For JA, the mobile phase was prepared by mixing methanol and 0.1% formic acid (65%:35%, v/v). Injection volume was 10 μL, flow rate 0.8 ml/min, column temperature 35°C, aliasing time 30 min and monitored at 230 nm. At least three independent biological replicates of each sample were performed.

### Protein Preparation

Leaves tissue (approximately 0.1 g FW for each biological replicate) was ground into a fine powder in liquid nitrogen and thoroughly transferred to an Eppendorf tube. To the tube, 1 ml of pre-cooled phenol extraction buffer was added, incubated the mixture at room temperature for 10 min and then added 1 ml phenol saturated with Tris-HCl (pH 8.0). The mixture was shaken for 40 min at 4°C. After centrifugation at 12,000 g for 15 min at 4°C, the upper phenolic phase was then collected, the debris was removed and the protein was precipitated with pre-cooled 100 mM ammonium acetate-methanol solution for 12 h at −20°C. After centrifugation, the pellet was washed three times with cold acetone and air-dried for 5 min. The protein was resuspended in 600 μL SDT (4% SDS, 100 mM Tris-HCl, 1 mM DTT, pH7.6), boiled water bath for 5 min and centrifuged at 12,000 g for 10 min at room temperature. The sample was collected and stored at −80°C for iTRAQ analysis. The protein concentration was determined according to Bradford assay (Bradford, 1976).

### Trypsin Digestion and iTRAQ Labeling

Approximately 100 μg protein of each biological replicate was used for digestion. Firstly, the protein sample was reduced by the addition of 120 μL buffer (10 mM DTT, 8 M Urea, 100 mM TEAB, pH 8.0) at 60°C for 1 h, and then alkylated using 50 mM iodoacetamide for 40 min at room temperature in the dark. Subsequently, the protein sample was diluted with 100 μL 100 mM TEAB. Then, 2 μL sequencing-grade trypsin (1 μg/μL; Promega) was added for the digestion at 37°C for 12 h. After centrifugation at 12,000 g for 20 min, the supernatant was collected and lyophilized. Briefly, one unit of the iTRAQ reagent (defined as the amount of reagent required to label 100 μg of protein). The sample was thawed and reconstituted in 100 μL 100 mM TEAB. The 100 μL iTRAQ reagent was transferred to the sample tube and labeled differently by incubation for 2 h at room temperature. After addition of 200 µL water to quench the labeling reaction, the solution was lyophilized.

### Mass Spectrometry Analysis

The labeled peptides were subjected to nanospray Flex source and analyzed by Q-Exactive mass spectrometer (Thermo, USA). Samples were loaded by a capillary C18 trap column (2 cm× 75 µm) and then separated by a C18 column (15 cm×75 µm) on an EASY-nLC™ 1200 system (Thermo, USA). The flow rate was 300 nL/min and linear gradient was 90 min (from 5 - 85% B over 67 min; mobile phase A= 2% ACN/0.1% FA and B=95% ACN/0.1% FA). Full MS scans were acquired in the mass range of 300-1800 m/z with a mass resolution of 70000 and the automatic gain control (AGC) target value was set at 1000000. The twelve most intense peaks in MS were fragmented with higher-energy collisional dissociation (HCD) with a normalized collision energy of 28%. MS/MS spectra were obtained with the tandem mass spectrum resolution of 35000, an AGC target of 50000 and a max injection time of 100 ms. The Q-E dynamic exclusion was set for 30.0 s and run under positive mode.

### Protein Identification and Quantification

The resulting MS/MS Spectral data files were analyzed using Proteome Discoverer™ 1.3 (Thermo, USA) software using the SEQUEST® search engine and searched against a Uniprot *Triticumaestivu*. fasta database for wheat at 1% false discovery rate (FDR). Mass error of precursor ions and fragment ions was set to 10 ppm and 0.02 Da, respectively.

The screening criteria of reliable proteins were as follows: unique peptide ≥1, removal of invalid values and antilibrary data, and screening of differentially expressed protein based on the reliable proteins. To screen differential proteins a Student’s *t*-test *P* < 0.05 and fold change > 1.3 or < 0.77 across the flag leaf samples or more than 1.5-fold or less than 0.67-fold were selected across the grain samples from a control and two treatments were applied based two experiments.

### Bioinformatics Analysis

The database (http://www.omicsbean.com:88/) and the OmicsBean software (http://www.ebi.ac.uk/interpro/) were used for gene ontology (GO) annotation. In the OmicsBean software, each protein was assigned to biological processes, cellular components and molecular functions. After annotation, proteins were mapped to the pathways in Kyoto Encyclopedia of Genes and Genomes (KEGG) database (http://www.kegg.jp/). Protein-protein interaction (PPI) analysis was performed using the Cytoscape software (Shannon *et al*., 2003), in which the threshold value (confidence cutoff) was set at 400, when the confidence score of the potential PPI was high, as indicated by solid lines or dashed lines.

### Enzyme Assays

For total amylase activity, flag leaf (approximately 0.1 g FW) was homogenized in a pre-chilled mortar and pestle with 1 ml cooled distilled water. After adding another 9 ml of distilled water, the mixture was placed at room temperature for 20 min to extract total amylase. Then the homogenate was centrifuged at 12,000 rpm for 15 min at 4°C. The supernatant was separated and used as the enzyme extract. To 1 ml enzyme extract, 1 ml soluble starch (1%, w/v) or 1 ml distilled water (control) was added and incubated at 40°C for 5 min. Subsequently, 2 ml of 3, 5-dinitrosalicylic acid reagent (1% (w/v) 3,5-dinitrosalicylic acid and 100 mM phosphate buffer (pH 7.0)) was added to the mixture before heating in a boiling water bath. Absorbance was measured at 540 nm using a spectrophotometer. To measure α-amylase (EC 3.2.1.1) activity, β-amylase (EC 3.2.1.2) in the crude enzyme extract was inactivated by heating at 70°C for 15 min. All of the other steps were the same as the assay of total amylase activity described above. The activity of β-amylase was calculated as the difference between total amylase and α-amylase activity. The enzyme activity was expresses as the amount of enzyme catalyzing to the production of 1 mg reducing sugar per min per mg protein.

Protease (EC 3.4.21.40) activity was determined spectrophotometrically using casein as the substrate. Flag leaf (approximately 0.1 g FW) was ground in a mortar. The homogenates were centrifuged at 12000 × g for 15 min at 4°C and the supernatant was transferred to a test tube for measurement of proteases activity. The reaction mixture containing 1 ml enzyme extract and 1 ml casein (2%) were heated at 40 °C for 10 min. Trichloroacetic acid solution (2 mL 5%) was added into the reaction for 20 min at 40°C. After centrifugation (12,000×g, 10 min), a reaction mixture containing 1 mL supernatant, 5 mL Na_2_CO_3_ (0.5 M), and 1 ml Folin reagent was prepared and its absorbance was then colorimetrically measured at 680 nm. The pH levels of acid proteinase, neutral proteinase, and alkaline proteinase buffer solutions were 3.6, 7.5 and 11, respectively. The activity of proteases was estimated by measuring residues of tyrosine and other aromatic amino acids released from hydrolyzed proteins according to a standard curve. The activity was expressed as nmol tyrosine released per min per mg protein.

### Grain Mass

Eighty plants were harvested from each plot at maturity and threshed by hand. The grains were oven-dried at 60°C to a constant weight. Grain mass were then weighed and grain number were counted to obtain single grain mass. Grain mass and grain number were showed based on half-spike.

## Acknowledgements

This work was supported by the Natural Science Foundation of Shandong Province (ZR2016CM39), the Shandong Project of Leading Talent for Mount Tai Industry (LJNY201601), the Shandong Modern Agricultural Technology and Industry System (SDAIT-01-06) and the National Earmarked Fund for Modern Agro-industry Technology Research System (CARS-3-1-21). We Thank Shanghai Luming Biotechnology CO., LTD for proteomics services and bioinformatic support.

